# FactorialHMM: Fast and exact inference in factorial hidden Markov models

**DOI:** 10.1101/383380

**Authors:** Regev Schweiger, Yaniv Erlich, Shai Carmi

## Abstract

**Motivation:** Hidden Markov models (HMMs) are powerful tools for modeling processes along the genome. In a standard genomic HMM, observations are drawn, at each genomic position, from a distribution whose parameters depend on a hidden state; the hidden states evolve along the genome as a Markov chain. Often, the hidden state is the Cartesian product of multiple processes, each evolving independently along the genome. Inference in these so-called Factorial HMMs has a naïve running time that scales as the square of the number of possible states, which by itself increases exponentially with the number of subchains; such a running time scaling is impractical for many applications. While faster algorithms exist, there is no available implementation suitable for developing bioinformatics applications.

**Results:** We developed FactorialHMM, a Python package for fast exact inference in Factorial HMMs. Our package allows simulating either directly from the model or from the posterior distribution of states given the observations. Additionally, we allow the inference of all key quantities related to HMMs: (1) the (Viterbi) sequence of states with the highest posterior probability; (2) the likelihood of the data; and (3) the posterior probability (given all observations) of the marginal and pairwise state probabilities. The running time and space requirement of all procedures is linearithmic in the number of possible states. Our package is highly modular, providing the user with maximal flexibility for developing downstream applications.

**Availability:** https://github.com/regevs/factorialhmm

## 1 Introduction

Hidden Markov models (HMMs) are instrumental for modeling sequential data across numerous disciplines, such as signal processing, speech recognition, and climate modeling. HMMs are also widely popular in bioinformatics (Durbin *et al*., 1998; Ernst and Kellis, 2012; Li *et al*., 2014; Shihab *et al*.), due to the sequential nature of the genome. In a standard HMM, the system is characterized by an internal discrete state variable, which evolves as a Markov chain between time points. In typical applications in bioinformatics, the state describes a slowly varying characteristic of the genome that is not directly observed, such as the identity or population of origin of the ancestor, or the chromatin state. Under the model, observed data points (which may be continuous or discrete) are drawn, at each genomic position, from a distribution whose parameters depend on the hidden state at that position. Algorithms exist for the efficient inference, given a sequence of observations, of the most likely sequence of hidden states or the marginal probability distribution of the state at each position, among other quantities.

An important generalization of the HMM is the factorial HMM (Ghahramani and Jordan, 1997), in which there are multiple independent Markov chains of latent variables, and the distribution of the observed variable at a given time step is conditional on the states of all of the latent variables at that same time step. Factorial HMMs have been particularly useful in genetics, where they have been used in classic linkage algorithms (Lander and Green, 1987), admixture mapping (McKeigue *et al*., 2013), local and global ancestry inference (Bercovici *et al*., 2012; Baran *et al*., 2012; Pei *et al*., 2018), estimating identity-by-descent (IBD) (Bercovici *et al*., 2010), identifying relationships (Kyriazopoulou-Panagiotopoulou *et al*., 2011), and detecting recombination events (Husmeier, 2005).

In a general implementation of an HMM, the time and space complexity is quadratic in the number of states. For a factorial HMM, the number of states is exponential in the number of latent Markov chains. However, the independence of the hidden chains in the factorial HMM can lead to reduced complexity of several standard operations. In (Ghahramani and Jordan, 1997), an exact calculation is presented to perform the Forward-Backward algorithm in *O*(*TMK^M+1^*), instead of *O*(*TK*^2M^), where *T* is the number of time steps (or genomic positions), *K* is the alphabet size of each latent chain, and *M* is the number of latent chains.

We extended the Forward-Backward efficient calculations presented in (Ghahramani and Jordan, 1997) to perform fast exact calculation of all standard HMM operations, effectively avoiding both time and space quadratic complexity. We present our implementation of these methods in FactorialHMM, a Python program supporting the full set of standard HMM operations. FactorialHMM is expected to be particularly useful in the context of genetics, providing a flexible and generic framework for researchers.

## 2 Features

We highlight the main distinct features of FactorialHMM. First, the latent Markov chains may be inhomogeneous, i.e., a distinct transition matrix per chain and per step. This is an important feature in genetics applications, where the (physical or genetic) distance between consecutive genomic sites may vary. Second, the alphabet size of each latent chain may be distinct, which allows for flexibility of modeling. The observed states may be continuous or discrete, and univariate or multivariate. Finally, the library supports exact and efficient calculations of all standard HMM operations, as follows. (i) Simulation from the model. (ii) Calculation of the marginal posterior probabilities of the latent hidden states, using the Forward-Backward algorithm. (iii) Calculation of the maximum posterior sequence of hidden states given the observed sequence, using the Viterbi algorithm. (iv) Calculation of the posterior pairwise probability for the hidden states, per chain, conditional on the observed data, used for constructing an efficient M-step in the Expectation-Maximization (EM) algorithm. (v) Sampling from the posterior distribution of hidden states, again given the observed sequence. The full description of these methods and the details on their efficient implementation are given in the Supplementary information.

## 3 Performance

We are not aware of an available stable implementation of the factorial HMM to which to compare, although several repositories contain some support for it (Johnson and Willsky, 2013; Ghahramani and Jordan, 1997). To demonstrate the computational efficiency of FactorialHMM, we compared it to a naïve implementation of the HMM using *hmms*, whose core algorithms are implemented in Cython. We tested a factorial HMM with *M* chains with binary states, with a symmetric transition matrix, uniform initial state distributions, and 100 observations simulated from the HMM with random initial states. We measured the computation time for the Viterbi and the Forward algorithms for increasing values of *M*. As expected, the computation time grows much faster for the naïve HMM than for FactorialHMM, allowing the user to scale up the number of hidden states.

**Figure 1.**
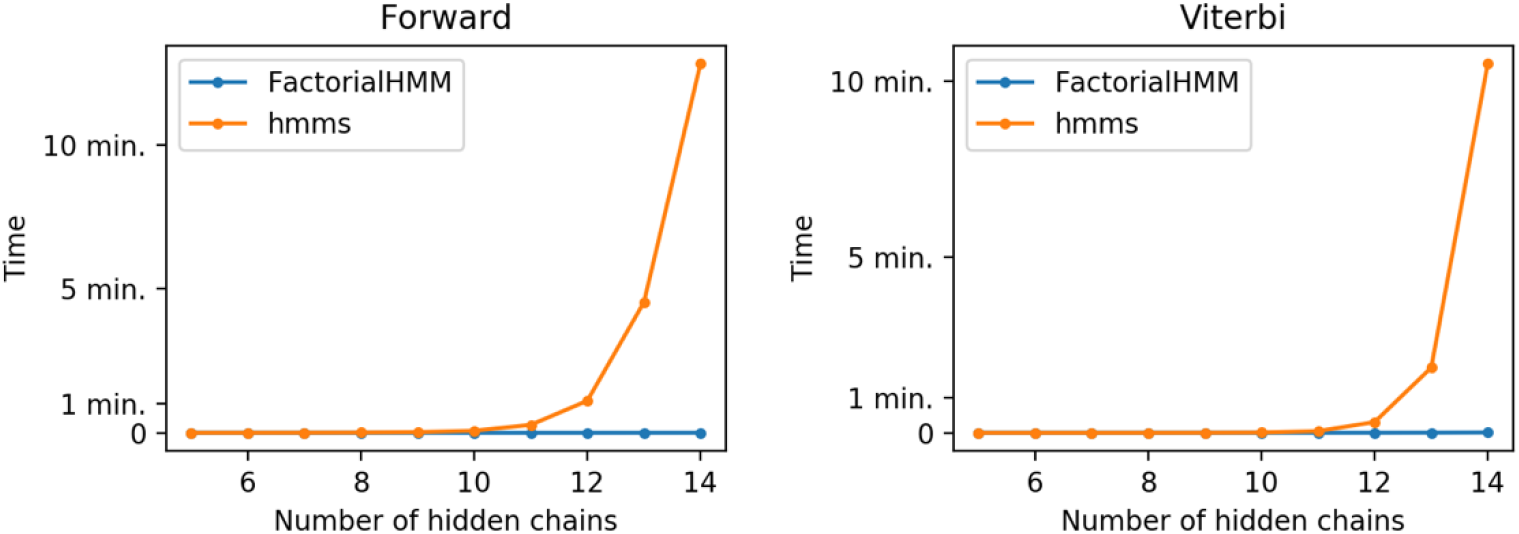
A benchmark comparing FactorialHMM and a naïve implementation by hmms. We tested a factorial HMM with M chains with binary states and 100 observations simulated from the HMM with random initial states. We compared the performance of both implementations on the Forward and the Viterbi algorithms. The computation time grows much faster for the naïve HMM than for FactorialHMM. All computation times of FactorialHMM were <1sec.

## 4 Discussion

Future directions for development include: (i) implementing the EM for specific forms of the transition and emission matrices; (ii) low-level language implementation; and (iii) an implementation of the linear-time approximate inference procedures detailed by (Ghahramani and Jordan, 1997). We look forward to comments, suggestions and future collaborative development of FactorialHMM.

Factorial HMM is available at https://github.com/regevs/factorial_hmm.

